# Prevalence and risk factors for avian influenza in backyard pigeons, ducks, and chickens in District Toba Tek Singh Pakistan

**DOI:** 10.1101/2024.11.08.622629

**Authors:** Iram Shakeel, Hamad Bin Rashid, Quratul Ain, Alijaan Inayat, Umer Shakeel, Adedayo Michael Awoniyi, Mamoona Chaudhry

## Abstract

Avian influenza virus (AIV) is a zoonotic disease that can be transferred from birds to humans. Various pandemics of AIV have drastically affected the poultry industry and backyard birds (ducks, chickens, and pigeons) worldwide, such as in Europe, the USA, Africa, and Asia. In Pakistan, numerous outbreaks of H7, H5, and H9 in poultry industries have been documented in commercial and rural areas, resulting in financial losses. The epidemiological status of different AIV subtypes in backyard birds in rural areas is largely unknown. Cross-sectional study was conducted in 2018-2019 with multistage clustering sampling. These tracheal and cloacal pooled swab samples were collected and tested for AIV. Positive AIV pooled swab samples were further evaluated at the individual level. RNA was extracted using the Trizol method followed by multiplex RT‒PCR with primers and probes to identify the M-gene and subtypes of the AI virus. Statistical analysis was performed by using the multivariate logistic regression model. Overall, the prevalence of AIV in backyard chickens, pigeons, and ducks was 13.0%, 7.7%, and 11.3%, respectively. Among AIV subtypes, H7, H9, and HA/Untyped commonly circulated within backyard birds. All cities had nonsignificant differences (*p* > 0.05) in the prevalence of AIV among pigeons, ducks, and chickens. According to univariate logistic regression analysis, 16 of the 32 risk factors were further selected for the multivariate logistic regression model. Two potential factors were identified to be significantly associated (*p* < or =0.05). Among the variables, species (pigeon OR 9.2, 95% CI: 1.92-44.2; *p*=0.006) and chicken (OR 16.2, 95% CI: 2.89-91.5; *p*=0.006) strongly associated with the incidence of AIV among cities. Another variable, fighting cock, had a moderate association (OR 2.25, 95% CI: 0.99-5.09; *p*=0.051) with AIV prevalence. Enhancing surveillance, biosecurity, educational intervention, and farm practices to prevent AIV and improve poultry production and public health in Pakistan.

## Introduction

Avian influenza virus (AIV) is a zoonotic disease that is often spread in animals, chiefly birds, and sometimes to humans (1). Four types of influenza viruses exist, A, B, C, and D, with only type A generally capable of causing global pandemics in humans (2). Influenza A viruses belong to the family Orthomyxoviridae and genus influenza virus, and are divided into subtypes based on the stereotyping of eighteen hemagglutinin (H1-18) and eleven neuraminidase (N1-11) genes (3), with the H5 and H7 responsible for most infections ranging from asymptomatic to mild with high mortality rates, especially in cases of co-infection (4). AIV consists of two pathotypes, including low pathogenic avian influenza (LPAI) and high pathogenic avian influenza (HPAI) representing viruses that cause mild and severe disease and death rates in poultry (5).

For the past two to three decades, AIV has resulted in a global loss in the poultry industry amounting to millions of USD and significant human infections and deaths, particularly in many Low-and Middle-Income countries in Asia and Africa (6). For example, evidence of endemic H9N2 virus has been reported among commercial and backyard poultry in Asia and Europe (7–9). In most Asian countries, AIV seems to impact all sectors of the poultry population, with the first highly pathogenic AIV (H7N3) notably reported in Pakistan in 1994, which eventually resulted in approximately 3 million birds amounting to more than 6 million USD losses in the country’s agricultural sector (10). Humans may become infected with influenza virus either through direct contact with infected birds/animals during slaughtering or processing, or indirectly through contact with contaminated environments such as drinking water and agricultural resources that are contaminated by the secretions of infected birds (2).

The continued increase in the number of human populations in many LMICs countries has necessitated the rapid expansion in the agricultural sector to meet both human protein and employment needs (11). For example, poultry is a growing portion of the agricultural sector in Pakistan, annually experiencing approximately 10% growth rate and yielding close to 20 billion eggs to meet the protein needs of Pakistanis (10). As a result, backyard poultry is the main source of livelihood in many rural areas in the country, albeit with poor biosecurity measures, leading to potential risks of AIV spillover to humans from animals (12). Despite recent evidence suggesting that ducks and pigeons could potentially contribute to the spread of AIV in the environment, with surveillance conducted in Iran in 2016 during the H5N8 outbreak confirming the presence of AIV in commercial and backyard poultry, including wild birds (13), and another study confirmed H5 endemic among ducks (2.4%) in Egypt suggesting the need to develop specific prevention measures against future outbreak (14).

However, few or no studies have estimated the prevalence of AIV among domesticated (particularly, pigeons, ducks, and chickens) and the associated risk factors in Pakistan despite reports of several outbreaks of H7, H5, and H9 reported in poultry industries, leading to significant economic losses (15, 16). Different risk factors may be associated with the transmission of avian influenza, including contact with infected domestic birds or environment, distance from poultry, previous disease history, and contact with wild birds (16, 17). Given that most Pakistanis reside in rural areas, with almost every home possessing backyards where they raise birds, either for personal or commercial consumption (16), as a result, increasing the risk of AIV spillover to humans, this study aimed to evaluate the prevalence and pattern of AIV transmission, and the associated risk factors encouraging the spread of the virus among backyard poultry (pigeons, ducks, and chickens) in the District of Toba Tek Singh, Pakistan. Results from this study will be useful in guiding against future outbreaks, especially in poor communities lacking satisfactory biosecurity measures.

## Material and Methods

### 2.1: Study Area

This study was conducted in the District Toba Tek Sigh, which is located in the Centre of Punjab, Pakistan. It is bounded by Faisalabad to the east, Jhang to the west, and the Ravi River to the north. It is located between 30°33’ and 31°2’ degrees north latitude and between 72°08’ and 72°48’ degrees longitude. District Toba Tek Singh has four Tehsils, Gojra, Kamalia, Toba Tek Singh, and Pir Mahal. The population of district Toba Tek Singh is 2,191,495 (18). The majority of income sources are associated with agriculture, livestock, and fishing. Toba Tek Singh is 2^nd^ largest hub for poultry farming in Pakistan (19).

### 2.2: Sample Design

We employed multistage clustering sampling during the process. We selected district Toba Tek Singh and 4 clusters (Toba Tek Singh, Gojra, Kamalia, and Pir Mahal) as non-randomly selected cities within it. Probability proportional to size with replacement was used to select 30 villages as primary sampling units randomly, using C-Survey, version 2.0 to calculate sample size (20). This method was chosen as there was no complete sampling frame available for the population. Finally, eight households were systemically chosen from each village for bird sampling using the spinning pencil method for sampling (21).

### 2.3: Sample Collection

We conducted a cross-sectional sampling between 2018 and 2019. A total of 240 samples (tracheal/oropharyngeal and cloacal samples) were collected from healthy backyard birds with 5 tracheal and 3 cloacal pooled swab samples, and the positive pools were assessed at the individual level. The backyard birds included ducks, pigeons, and chickens from villages in district of Toba Tek Singh. The samples were collected from live birds by restraining them. A sterilized swab was used to collect samples from each bird. Swab samples were transferred into brain heart infusion media supplemented with antibiotics in marked cryovials. The ice packs were placed in an icebox to maintain the cold chain temperature during transportation. A complete predesigned questionnaire was completed after the owners provided their consent. The questionnaire contained information regarding sociodemographic and poultry-related questions/variables such as age, previous disease signs, distance from commercial poultry farms, contact with wild birds, equine, dog, and feline, morality and recovery history, feed history, rearing places, commercial poultry farms-related job history in the family, medication, and vaccination history, and water source.

### 2.4: Virus Propagation

All samples were aggregated into tracheal samples (chickens) and cloacal samples (pigeons and ducks) and pooled separately. For virus propagation, the pooled samples were inoculated into 9 days of embryonated eggs and incubated at 37°C for 24-72 h. Fertilized eggs were incubated at 37°C and 50-60% humidity and monitored for 9 days for embryo development by an egg candle. For stock of virus, after incubation, the allantoic fluid was harvested from each egg and stored at -80°C after centrifugation at 12000 rpm for 10 min at 4°C (22).

### 2.5: RNA Extraction Method

RNA was extracted from allantoic fluid samples via the Trizol method. Then, 250 µL of fluid was mixed with 750 µL of Trizol agent by homogenization. The mixture was then mixed with 250 µL (100%) of chloroform, vortexed, incubated at room temperature for 7 min, and centrifuged at 12000 rpm for 15 min at 4°C. Subsequently, 450 µL of the mixture was extracted from the upper layer, and the mixture was transferred to a new numbered vial. Then, 500 µL (100%) of isopropanol was added for 10 min at room temperature, and the mixture was centrifuged as described above. After washing with 75% ethanol, the RNA precipitate was collected by centrifugation at 10000 rpm for 8 min at 4°C, dissolved in 50 µL of RNase-free water, and stored following methods previously described by Brauer et al.,(23).

### 2.6: RT‒PCR Method

cDNA was synthesized by extracting RNA from the sample with the help of a commercially available cDNA synthesis kit® **(Thermo Fisher Scientific, Lithuania)** following the manufacturer’s instructions. The following probes and primers were used for the detection of the M-gene of the AI virus, as described in **Table 1** following methods previously described by Rashid et al., (24). An RT‒PCR Kit® was used for the amplification of DNA via RT‒PCR. The 25 µl mixture contained 12.5 µl of PCR Master Mix, 2 µl of template, 1 µl of forward primer, 1 µl of reverse primer, and 8.5 µl of DEPC. The M-gene was amplified by RT‒PCR under the following conditions: a holding stage at 95°C for 10 min, a cycling stage at 95°C for 15 s and 95°C for 30 s, and a holding stage at 4°C at infinity. For cross-checking, each pool was subjected to positive and negative controls.

**Table 1.**
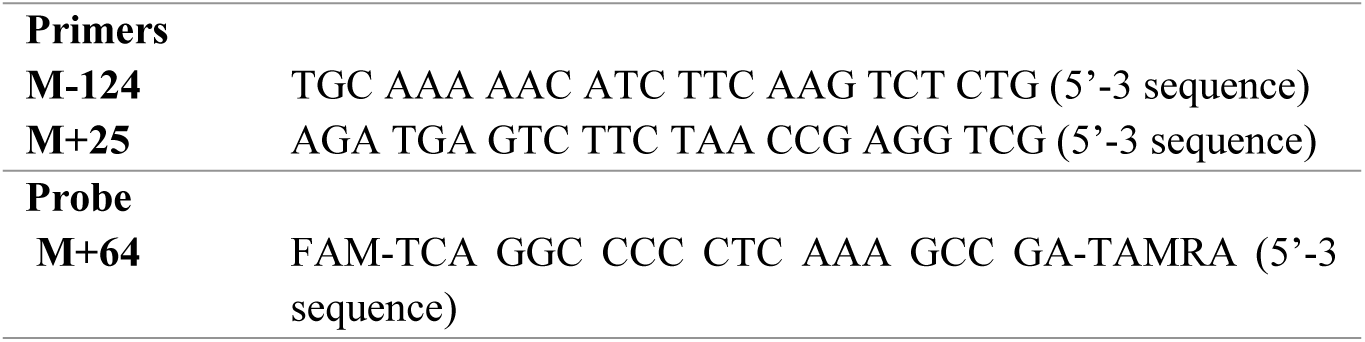
List of Probes and Primer of Avian Influenza Virus M-gene.

### 2.7: Ethical considerations

Approval was obtained from the Department Review Committee and Ethical Review Committee of the University of Veterinary and Animal Science Lahore. Informed consent was also obtained from the respondents, and the confidentiality of the data was strictly maintained.

### 2.8: Statistical analysis

Statistical analysis was performed by using Stata 16. All the data were documented on an Excel sheet and exported into Stata software for statistical analysis. We calculated point estimates of weighted prevalence with 95% Cl in backyard birds within each city. We used logistic regression with binary response (positive or negative AIV). At the initial stage, 32 exploratory variables were screened by univariate analysis. Out of 32 variables, 16 risk factors fulfilled the selection criterion (Wald test *p*<0.25) and were further subjected to multivariate logistic regression. Before the multivariate logistic regression model, a collinearity test was applied between selected variables. To construct the final model, a stepwise model-building approach was used. The odds ratio and Wald test *p-value* equal to 0.05 for each variable were used to retain or remove selected variables. The final logistic model assessed an odds ratio (OR) with 95% Cl for each variable.

## Results

### Prevalence of AIV in backyard birds

Overall, we observed 13.0% prevalence of AIV in backyard chickens, with Pir Mahal having non significantly higher prevalence (15.3%) and Gojra similarly reporting a non-significant lower prevalence (11.5%) in the sampled cities thus indicating a uniform distribution of AIV prevalence among chickens across the four sampled cities (**Table 2**). Among the subtypes, the overall prevalence of H9 and H7 among backyard chickens was 10.2% and 2.80%, respectively.

**Table 2.** Positive tracheal samples of chicken for Avian influenza virus and its subtypes (H7and H9) by using qRT‒PCR in district Toba Tek Singh*.

| Cities | Total Sample<br>N | Subtype H9<br>n (%) | Subtype H7<br>n (%) | Positive AIV<br>n (%) |
| --- | --- | --- | --- | --- |
| Gojra | 26 | 3 (11.5) | 1 (33) | 2 (67) |
| Toba Tek Singh | 27 | 4 (14.81) | 0 (0) | 4 (100) |
| Kamalia | 25 | 3 (12) | 1 (33) | 2 (67) |
| Pir Mahal | 26 | 4 (15.3) | 1 (25) | 3 (75) |
| **Total** | 107 | 14 (13.0) | 3 (21) | 11(79) |

\*There were no significant differences between the prevalence of Avian influenza virus and Cities among chickens (*p-value=0.98*)

In general, we reported 11.3%, prevalence of AIV in backyard ducks, with a non-significant high prevalence (11.1%) observed in Toba Tek Singh and a low prevalence of (7.5%) reported in Gojra in the sampled cities (**Table 3**). Among the subtypes, the overall prevalence of H7 and HA/Untyped among backyard ducks was 6.8% and 4.54%, respectively. Positive samples that were not positive for AIV subtypes H5, H9, and H7 were labeled as AIV HA/Untyped.

**Table 3.** Positive cloacal samples of duck for Avian influenza virus and its subtypes (and HA/Untyped) by using qRT‒PCR in district Toba Tek Singh*.

| Cities | Total Sample<br>N | Positive AIV<br>n (%) | Subtype H7<br>n (%) | HA/Untyped<br>n (%) |
| --- | --- | --- | --- | --- |
| Gojra | 15 | 2 (7.5) | 1 (50) | 1 (50) |
| Toba Tek Singh | 9 | 1 (11.1) | 0 (0) | 1 (100) |
| Kamalia | 10 | 1 (10) | 1 (100) | 0 (0) |
| Pir Mahal | 10 | 1 (10) | 0 (0) | 1 (100) |
| **Total** | **44** | **5 (7.69)** | **2 (40)** | **3 (60)** |

\*There were no significant differences between the prevalence of Avian influenza virus and Cities among ducks (*p*-value=0.99)

The overall prevalence of AIV in backyard pigeons was 7.7%, with a high non-significant prevalence (9.09%) reported in Toba Tek Singh and Pir Mahal and low prevalence (4.54%) in Gojra (**Table 4**). Among the subtypes, the overall prevalence of H7 and HA/Untyped among the backyard pigeons was 6.59% and 1.09%, respectively.

**Table 4:** Positive cloacal samples of pigeons for Avian influenza virus and its subtypes (H7 and HA/Untyped) by using qRT‒PCR in district Toba Tek Singh*.

| Cities | Total Sample<br>N | Positive AIV<br>n (%) | Subtype H7<br>n (%) | HA/Untyped<br>n (%) |
| --- | --- | --- | --- | --- |
| Gojra | 22 | 1 (4.54) | 1 (100) | 0 (0) |
| Toba Tek Singh | 22 | 2 (9.09) | 2 (100) | 0 (0) |
| Kamalia | 25 | 2(8) | 1 (50) | 1 (50) |
| Pir Mahal | 22 | 2(9.09) | 2 (100) | 0 (0) |
| **Total** | **91** | **7 (7.69)** | **6 (6.59)** | **1 (1.09)** |

\*There were no significant differences between the prevalence of Avian influenza virus and Cities among ducks (*p*-value=0.97)

Figure 1 and figure 2 show positive samples of AIV and its subtypes (H7, H9, and HA untyped) from ducks, pigeons, and hens in District Toba Tek Singh.

**Figure 1:**
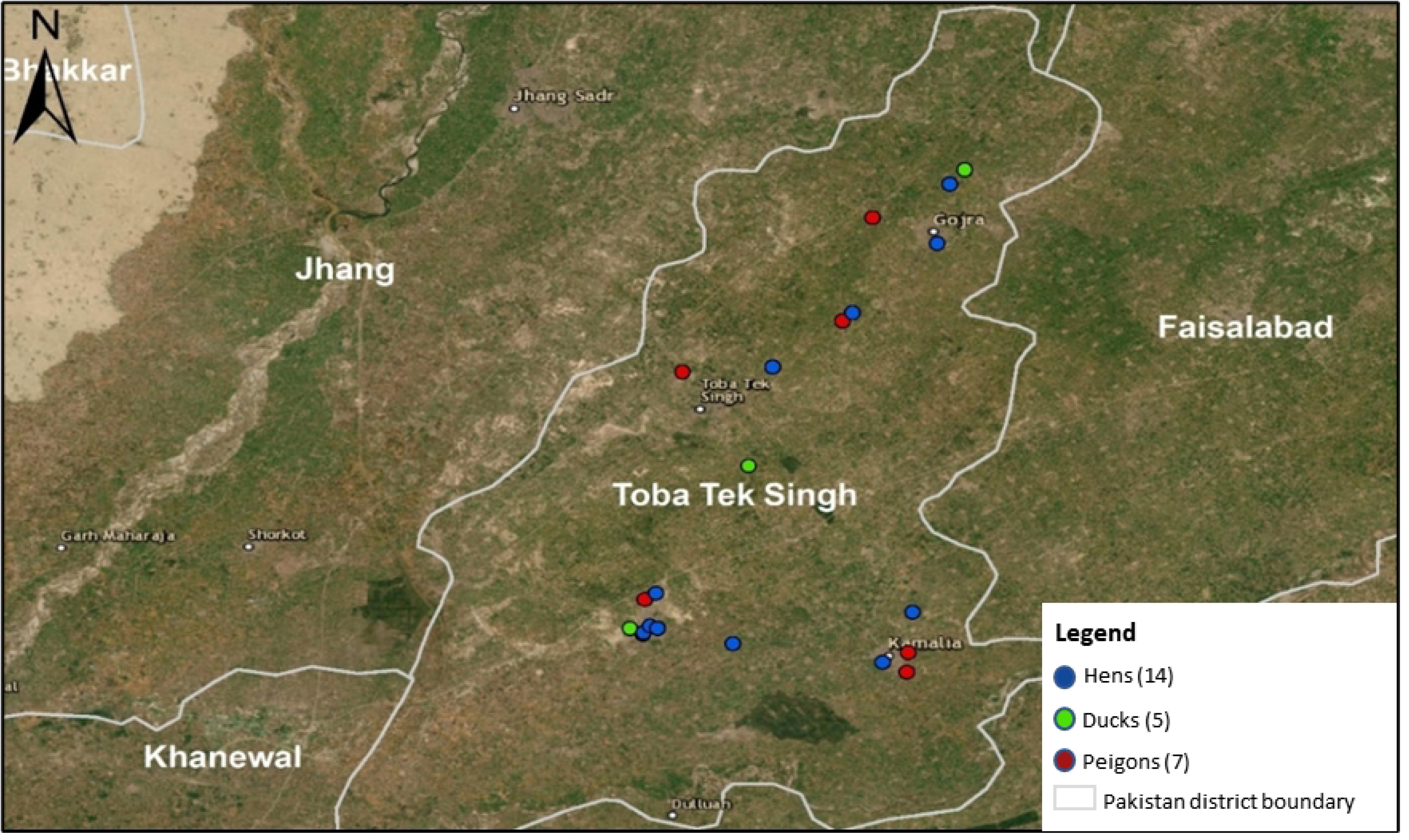
Positive samples of AIV in ducks, pigeons, and hens in the District Toba Tek Singh.

**Figure 2:**
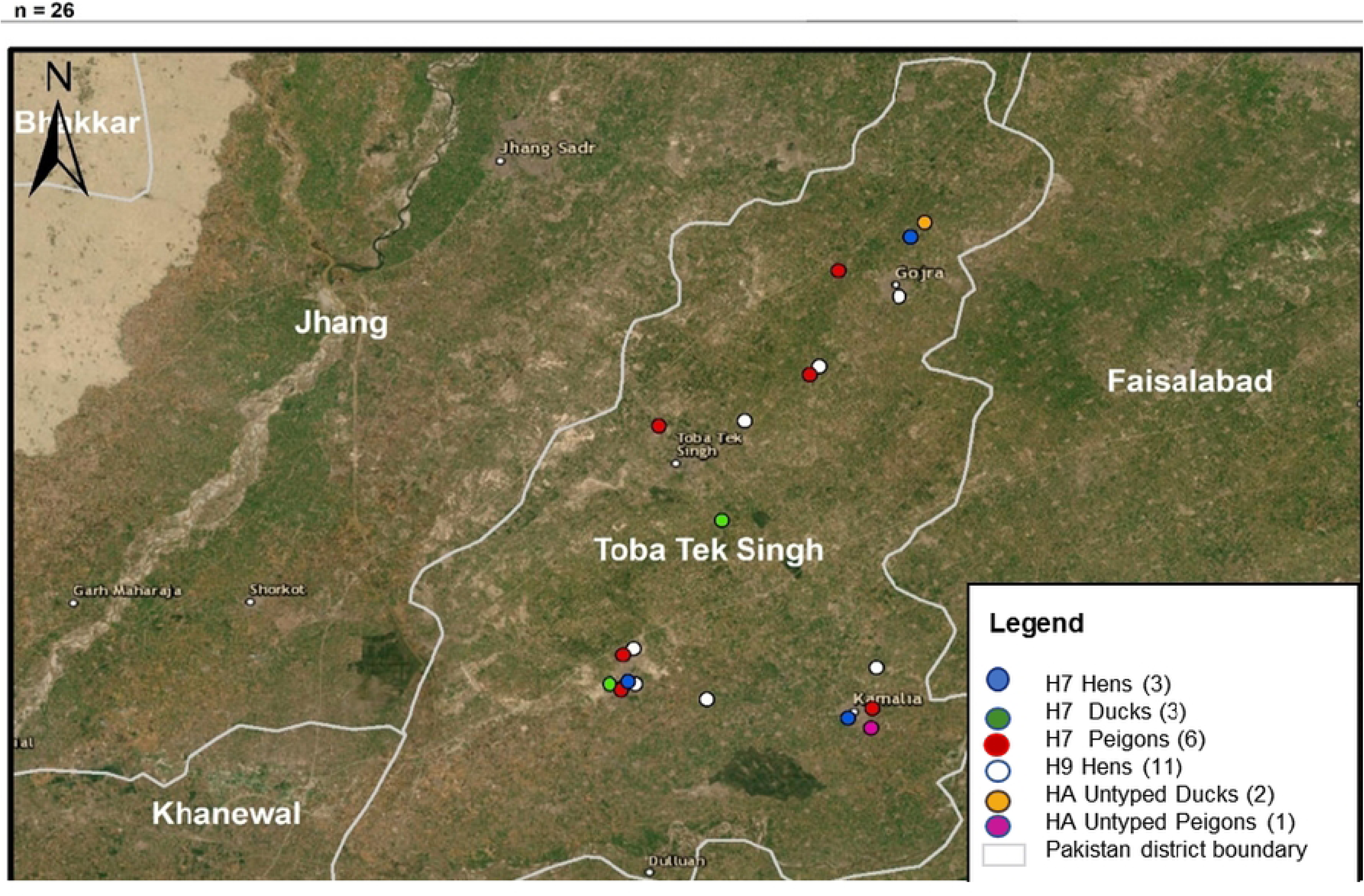
Positive samples of H7, H9, and HA-Untyped AIV subtypes in ducks, hens, and pigeons in the district of Toba Tek Singh.

### Risk factors associated with the prevalence of AIVs

Initially, 46 variables were selected for univariate logistic regression, but due to their association with the study outcome (birds positive or negative for AIV), 32 variables (risk factors) were included in the univariable logistic regression. Out of the 32 variables, 16 risk factors fulfilled the selection criterion (Wald test *p*<0.25). They were further included in the multivariable logistic regression model **(Table in S1 Table**) in the supporting information file named **S1 Table**.

According to the final model, two potential risk factors were significantly associated with the prevalence of AVI and its subtypes in backyard birds (p < or =0.05). Among the variables, species (pigeon OR 9.2, 95% CI: 1.92-44.2; p=0.006) and (chicken OR 16.2, 95% CI: 2.89-91.5; p=0.006) were strongly associated with the incidence of AIV among cities. Another variable, fighting cock, had a moderate association (OR 2.25, 95% CI: 0.99-5.09; p=0.051) with AIV prevalence (**Table 5**).

**Table 5.** Results of multivariant analyses of risk factors for AIV prevalence in backyard birds in District Toba Tek Singh.

| Risk Factors | Response Level | Positive Samples | Odds Ratio (OR) | CI (95%) | *p*-value |
| --- | --- | --- | --- | --- | --- |
| **Birds** | 242 | 104 |  |  | **0.006** |
| Duck | 44 | 9 | Ref |  |  |
| Pigeon | 91 | 39 | 9.21 | 1.92-44.2 |  |
| Chicken | 107 | 56 | 16.2 | 2.89-91.5 |  |
| **Fighting Cock** | 242 | 104 |  |  |  |
| No | 65 | 21 | Ref |  |  |
| Yes | 177 | 83 | 2.25 | 0.99-5.09 | **0.051\*** |

\*There were moderate significant differences in prevalence between the prevalence of Avian influenza virus and fighting cock in multivariate analyses (**p-value=0.051**).

## Discussion

The current expansion of the Pakistani population leading to higher demand for protein and intense competition for resources has resulted in most urban, peri-urban, and rural residents heavily investing in poultry farming particularly in their backyards and peri-domestic areas with poor biosecurity measures, eventually leading to significant mortality in poultry sector and high probable of virus spillover from the birds to humans. Surveillance of Avian influenza (AI) viruses in backyard poultry is a good early warning system for controlling outbreaks. In this study, we estimated the prevalence of AIV and its subtypes in domestic birds, namely ducks, pigeons, and chickens. The overall AIV prevalence in backyard chickens was 13.0% with subtypes H9 (10.2%) and H7 (2.80%) prevalence respectively in district Toba Tek Singh. Our results were similar to the prevalence of H9 (9.2%) and H7 (1.9%) previously reported in India (25), further supporting the claim that AIV is endemic to Asian countries. However, our results are lower than the high prevalence rates reported in Chile (26) and Vietnam (27), which could be attributed to variations in sampling strategies, study populations, and small sample sizes.

The overall AIV prevalence in backyard pigeons and ducks was 11.3% and 7.69% in our study, which is similar to previous results from China with H7 and HA/untyped AI subtypes (28). Our results differ from other findings in Kosice (29) and Egypt (30), which reported higher prevalence rates, which could be due to differences in testing techniques, population size, and geographic distribution. Identification of HA-untyped AI subtypes revealed the presence of other subtypes, such as H3N6 among backyard poultry, likely due to interactions with migratory birds near villages. Non-detection of H5 in this study might be due to features such as seasonal variation, migratory patterns, and ecological conditions with diverse geographic limitations (31).

Among the tehsils in district of Toba Tek Singh, Toba Tek Singh has a higher prevalence than other cities. This difference might be attributed to the dense population and city, poultry farming practices variation such as free-range, farming, conventional farming, and semi-cage or cage-free farming, and prominent free trading and transportation in this city compared to other cities. The prevalence of AIV in this study could be responsible for the previously reported high prevalence of AIV among poultry workers in Toba Tek Singh (32).

For the past two decades, backyard poultry has significantly contributed to the source of income of many rural families in Pakistan, but amidst poor biosecurity measures. This practice often predisposes backyard poultry farmers to AIV, in addition to farmer’s poor awareness or knowledge of AIV transmission mechanisms and how to best prevent its transmission. In this study, we identified fighting cocks as a significant risk factor (*p < 0.05*). Similarly, a study in Thailand revealed that fighting cocks are associated with the transmission of influenza. The regular movements of owners fighting cocks at the national and international levels may be associated with an increased risk of disease transmission. To effectively manage AIV transmission, intervention programs should encompass awareness/education campaigns, vaccination, and structured biosecurity systems at both the rural and peri-urban levels and should be regularly supported by policymakers (33). Similarly, proper monitoring of the purchasing and movement of fighting cocks should be encouraged to control the spread of infection in backyard poultry (34).

Species type was another significant risk factor (*P <0.006*), indicating that impoverished biosecurity in backyard poultry could lead to infection in these birds, as they have easy access to the outdoors, where they share the same feed and water sources as wild waterfowl birds (35). Chicken (OR 16.2, 95% CI: 2.89-91.5; *p<0.006*) was strongly associated with the prevalence of AIV among villages. This indicates that chickens could contribute to AIV transmission among humans, as they are kept in large farming systems, heavily dependent as sources of protein, and involved in global poultry trading, thereby contributing to the potential AIV transmission chain. Also, backyard poultry is easily scavenged by wild ducks, pigeons, and migratory birds near crops and water reservoirs in the process contributing to the expansion of AIV transition beyond just poultry and those having direct or indirect contact with poultry products (36). Therefore, monitoring and surveillance programs are crucial to manage transmission risk.

The association of cat presence with AIV infection in backyard poultry according to the result of our univariate analysis, might be associated with the circulation of HA/untyped AIV among backyard birds, which has been reported in literature, supporting the ability of cats to spread AI viruses among poultry and humans (37, 38). However, in another study, stray dogs had a stronger association with AI infection than cats (38). The presence of various animals near poultry farms, such as cats, pigs, and dogs, might possibly increase the risk of AIV transmission between diverse species. This finding might aid the prevention of secondary spreading, especially after an initial outbreak to reduce the widespread of AIV pandemics among animals and humans while also reducing economic loss. The association of history of medication with AIV infection in the results of our univariate analysis, with most (86%) poultry farmers preferring home remedies to medical remedies (39, 40) might predispose birds to AIV which is an alarming situation, and insist on the development of new inventions and increasing the number of veterinary staff at rural levels for awareness campaigns.

The authors recognized that there are several limitations and strengths in this study. For example, our small size somewhat prevents us from generalizing our results, the use of a cross-sectional study limits the ability to establish a causal relationship between the prevalence of AIV and risk factors, and due to financial constraints, sequencing of the untyped AIV subtypes was impossible. Also, however, our results proffer useful insight into the role of domestic birds in the circulation of AIV both among birds and the human community, and this information should be useful during intervention programs. Second, compared with serological tests, RT‒qPCR was used for the detection of AIV. Also, we suggest that more surveillance of the subtypes of AIV untyped HA should be conducted especially in backyard poultry to minimize zoonotic transmission in humans at the rural and peri-urban levels in Pakistan and other LMICs of similar profile.

## Conclusion

Backyard poultry have a great chance of becoming infected with AIV through contact with wild birds. Domestic ducks and pigeons can both spread infection with and without clinical signs and sources of H5, untyped HA, and H7. The high prevalence of this disease in domesticated birds indicates that this disease is endemic to rural and peri-urban areas making it a source of zoonotic disease. A surveillance system, including backyard poultry surveys, is needed, and based on these findings, control strategies should be encouraged by the Pakistani government. Control could be achieved by improving the biosecurity of backyard poultry, and changing farm practices such as controlled poultry housing systems or poultry farming systems. and educating farmers and poultry staff on the early detection of disease, and the biosecurity procedure useful for disease prevention. Backyard poultry remains a source of income among many Pakistanis, thus the policymakers should develop a program that involves all stakeholders to ensure that any proposed intervention is satisfactorily implemented to aid the country’s capacities to prevent, detect, and promptly respond to AIV.

## Acknowledgments

The authors are grateful for the contributions of participants in selected villages in district Toba Tek Singh and Punjab Livestock Department for providing data on backyard poultry. We are also grateful to all team members in the laboratory at the Department of Epidemiology and Public Health, University of Veterinary and Animal Sciences, Lahore.

## Conflicts of interest

All the authors declare no conflicts of interest.

## Authors contribution

**Conceptualization: Mamoona Chaudhry, Iram Shakeel, Hamad Bin Rashid**

**Data curation: Iram Shakeel, Umer Shakeel**

**Laboratory Analysis: Iram Shakeel, Mamoona Chaudhry, Quratul Ain**

**Data Analysis: Iram Shakeel, Mamoona Chaudhry, Alijaan**

**Project administration: Mamoona Chaudhry, Quratul Ain**

**Writing: Iram Shakeel, Adedayo Michael Awoniyi**

**Critical review: Adedayo Michael Awoniyi, Mamoona Chaudhry, Hamad Bin Rashid**

## Supporting information

**S1 Table. Univariate logistic regression analysis (DOCX)**

## Notes

### Competing Interest Statement

The authors have declared no competing interest.

